# Evolution of flowering time in a selfing annual plant: Roles of adaptation and genetic drift

**DOI:** 10.1101/2020.08.21.261230

**Authors:** Laurène Gay, Julien Dhinaut, Margaux Jullien, Renaud Vitalis, Miguel Navascués, Vincent Ranwez, Joëlle Ronfort

**Author notes:** **This article has been peer-reviewed and recommended by *Peer Community In Evolutionary Biology* (https://doi.org/10.24072/pci.evolbiol.100128)**.

## Abstract

Resurrection studies are a useful tool to measure how phenotypic traits have changed in populations through time. If these traits modifications correlate with the environmental changes that occurred during the time period, it suggests that the phenotypic changes could be a response to selection. Selfing, through its reduction of effective size, could challenge the ability of a population to adapt to environmental changes. Here, we used a resurrection study to test for adaptation in a selfing population of *Medicago truncatula*, by comparing the genetic composition and flowering times across 22 generations. We found evidence for evolution towards earlier flowering times by about two days and a peculiar genetic structure, typical of highly selfing populations, where some multilocus genotypes (MLGs) are persistent through time. We used the change in frequency of the MLGs through time as a multilocus fitness measure and built a selection gradient that suggests evolution towards earlier flowering times. Yet, a simulation model revealed that the observed change in flowering time could be explained by drift alone, provided the effective size of the population is small enough (<150). These analyses suffer from the difficulty to estimate the effective size in a highly selfing population, where effective recombination is severely reduced.

## Introduction

When facing changing environments, organisms can persist by one of three strategies: fleeing (migration), coping (plasticity) or adapting. If migration and plasticity can lead to rapid and reversible changes in the average phenotype of a population, adaptation proceeds through genetic changes and towards phenotypes with the highest fitness in a given environment. The literature describing adaptation in natural populations is vast (e.g. Bay et al., 2017; Côté and Reynolds, 2012; Kremer et al., 2012; Olson-Manning et al., 2012) and the recent rise of next generation sequencing has enabled tremendous progress in our knowledge about the genetic architecture of adaptation at the species level (Barrick and Lenski, 2013; Brown, 2012; Fournier-Level et al., 2011; Jones et al., 2012).

Long term temporal surveys (e.g. Visser, 2008), resurrection studies, where ancestors and descendants are compared under common conditions (see Box 1 in Franks, Weber, et al., 2014) or stratified propagule banks (Orsini et al., 2013) are powerful tools to reconstruct the evolutionary dynamics of populations that have faced environmental changes. Yet, observing a genetic change through time is not sufficient to claim that it is adaptation. Testing for selection as opposed to drift is one of the essential criteria for demonstrating adaptive responses, but is often overlooked (e.g. overlooked in 34% of the 44 reviewed studies based on phenotypic variation reviewed by Hansen et al., 2012). Demonstrating the influence of selection on a phenotypic change can be achieved by one of four methods (detailed in Table 2 in Hansen et al., 2012; Merilä and Hendry, 2014): reciprocal transplants (Blanquart et al., 2013), *Q*_ST_–*F*_ST_ comparisons (Le Corre and Kremer, 2012; Rhoné et al., 2010), genotypic selection estimates (Morrissey et al., 2012; Wilson et al., 2010), or tests of neutrality (pattern or rate tests, Lande, 1977). These methods all rely on measuring quantitative traits (fitness traits or traits supposed to be under selection) but require specific experimental settings. Pattern tests of neutrality rely on comparing evolution across replicates, for example by comparing phenotypic or allele frequency changes across replicates in experimental populations, or across natural populations, assuming that they are independent replicates of the evolutionary process (same effective size and selective pressure, no migration). Pattern tests can also apply through time if a long sequence of observations is available (Sheets and Mitchell, 2001). Alternatively, rate tests can be useful to examine the rate of genetic change in a population and compare it to the expectation under a neutral model with a given effective population size (Lande, 1976). The effective population size (thereafter *N*_e_) is defined as the size of an ideal Wright-Fisher population experiencing the same rate of genetic drift as the population under consideration (Crow and Kimura, 1970). Unlike experimental populations, where *N*_e_ can be monitored, an accurate estimate of *N*_e_ is required to perform such neutrality tests in natural populations. Temporal changes in allele frequency at neutral loci can be used to infer the effective size of the population considered (Nei and Tajima, 1981; Waples, 1989).

The ability for a population to adapt to environmental changes depends on several factors such as genetic variability, generation time, population size or mating patterns, in particular self-fertilization rates. In plants, a large fraction (40%) of species do, at least partially, reproduce through selfing (Goodwillie et al., 2005; Igic and Kohn, 2006). Selfing could challenge the process of adaptation because it directly decreases the effective population size *N*_e_ (reduced number of independent gametes sampled for reproduction (Pollak, 1987); increased homozygosity; reduced efficacy of recombination (Nordborg, 2000); increased hitchhiking and background selection (Gordo and B Charlesworth, 2001; Hedrick, 1980)). It is therefore expected that genetic variability is reduced in selfing populations, and empirical measures of diversity from molecular markers strongly support this prediction (Barrett and Husband, 1990; Glémin, Bazin, et al., 2006; Hamrick and Godt, 1996). Furthermore, several theoretical models also predict that selfing reduces quantitative genetic variation within populations (Abu Awad and Roze, 2018; D Charlesworth and B Charlesworth, 1995; Lande and Porcher, 2015), which has been recently confirmed by a meta-analysis of empirical data (Clo et al., 2019).

We can expect that this depleted genetic variation in predominantly selfing populations will limit their ability to adapt to changing environmental conditions and their long-term persistence and different theoretical models support this prediction (Glémin and Ronfort, 2013; Hartfield and Glémin, 2016; Kamran-Disfani and Agrawal, 2014). Yet, empirical data examining the response of predominantly selfing populations to environmental changes remain scarce, especially for data showing short term adaptation in the face of climate change (Qian et al., 2020). In a recent review focussing on evolutionary and plastic responses to climate change in plants, Franks, Weber, et al. (2014) reported “at least some evidence for evolutionary response to climate change […] in all of these studies”, and six of these 31 studies considered selfing populations.

Because there is no consensus between theoretical predictions, empirical and experimental data, the ability of selfing populations to adapt to environmental changes remains an open question. This calls for further fine scale population genetics analyses, with a focus on the evolutionary mechanisms involved and on the dynamics of adaptation. Here, we present a temporal survey in the barrel medic (*Medicago truncatula*) that enabled us to perform a resurrection study. *M. truncatula* is annual, diploid, predominantly self-fertilizing (>95% selfing, Bonnin, Ronfort, et al., 2001; Siol, Prosperi, et al., 2008) and has a circum-Mediterranean distribution. Flowering time is a major heritable trait (broad-sense heritability > 0.5, Bonnin, Prosperi, et al., 1997) that synchronizes the initiation of reproduction with favourable environmental conditions and could play a role in the adaptation to climate change. In *M. truncatula*, flowering time is highly variable along the distribution range and within some populations (Bonnin, Prosperi, et al., 1997). It is mainly controlled by two environmental cues: photoperiod and temperature (Hecht et al., 2005; Pierre et al., 2008). In the Mediterranean region, there has been a significant increase in temperatures between the 80s and nowadays accompanied by a decrease in mean precipitations (http://www.worldclim.org/). Most studies about adaptation in *M. truncatula* have so far relied on large collections of individuals representing the whole species with the aim of detecting selection footprints in the genome linked with flowering time (Burgarella et al., 2016; De Mita, Chantret, et al., 2011) or climatic gradients (Yoder et al., 2014). However, the complex population structure observed at the species level can make it difficult to understand the selective history of those genes (De Mita, Ronfort, et al., 2007). Indeed, natural populations of *M. truncatula* are composed of a set of highly differentiated genotypes that co-occur at variable frequencies (Bonnin, Ronfort, et al., 2001; Loridon et al., 2013; Siol, Prosperi, et al., 2008), a genetic structure typical for predominantly selfing species. How does this peculiar genetic composition constrain adaptation to changing environments remains unclear, but preliminary results in *M. truncatula* have shown that surveying the multilocus genotypic composition through time could reveal a large variance in the relative contributions of these genotypes to the next generations (Siol, Bonnin, et al., 2007). Here, we examined the temporal change of flowering time at the population level across 22 generations characterised by changing environmental conditions (temperature and rainfall). We describe the peculiar genetic structure of this highly selfing species and investigate the genetic mechanisms involved in adaptation. In particular, we test for the role of selection as opposed to genetic drift, following four steps. First, we consider the direction of the change in trait value in relation to the environmental change. Second, we estimate the extent of genotypic selection (Morrissey et al., 2012; Wilson et al., 2010) using selection gradients for flowering time based on several fitness estimates (including an estimate of the realised fitness based on changes in frequency of the multilocus genotypes through time). Then we estimate the effective population size, test the rate of evolution for neutrality by simulating how the frequency of the multilocus genotypes would change under genetic drift alone and explore the effect of the imprecision in the estimation of effective size. Finally, we examine the change in flowering time during the same time period at the regional scale, using one individual per population across the distribution range of *M. truncatula* in Corsica. A similar genetic change at the regional scale would lend weight to the hypothesis that the change in flowering time occurred in response to selection.

## Materials and Methods

### Studied population and experimental design

The focus population (F20089 or CO3 according to Jullien et al., 2019) is located in Cape Corsica (42°58.406’N - 9°22.015’E). In 1987 and 2009, around 100 pods were collected along three transects running across the population, with at least one meter distance between each pod collected, in order to avoid over-sampling the progeny of a single individual. Seeds collected in 1987 were stored in a cold room. In 2011, pods collected in 1987 and 2009 were threshed and seeds were replicated through selfing in standardized greenhouse conditions to control for maternal effects and build families of full-sibs produced by selfing. Seeds for this generation of multiplication were randomly selected from pooled samples of seeds from 1987 and 2009. 64 families collected in 1987 and 96 in 2009 were successfully multiplied. Out of these, 55 families for each of the two sampling years were randomly chosen in 2012. Seeds from the 110 families were scarified to ease germination and were transferred in Petri dishes with water at room temperature for six hours. We then used two different vernalization treatments (at 5°C during 7 or 14 days) to compare the vernalization requirement between the two years. Five replicates from each vernalization treatment were transferred back to the greenhouse, according to a randomized block design (five blocks and two treatments, adding up to a total of ten replicates per family, 1100 plants in total). Data loggers were placed on each table to monitor temperature and humidity. For each individual, the number of days after germination to form the first flower was recorded. In addition, the total number of seeds produced by each plant throughout its lifetime was measured as a proxy for fitness.

### Temporal changes in flowering time

Individual flowering times were converted to thermal times following Bonhomme (2000). The thermal time was calculated as the sum of the mean daily effective temperatures of each day between sowing and the emergence of the first flower, where the mean daily effective temperature is the day’s mean temperature minus the base temperature (*T_b_*). We used *T_b_* = 5°C, as reported by Moreau et al. (2007) for the *Medicago truncatula* reference line A17. Plants noted as sick or failing to produce leaves were removed from the data sets (22 individuals removed). Collected measures were tested for normality using quantile–quantile (Q-Q) plots (Nobre and Singer, 2007). All analyses were conducted using R version 2.15.2. We used linear mixed models (lme4 package) to test for a significant change in flowering time between the sampling years. The model included two fixed effects: sampling year (1987 or 2009) and treatment (short or long vernalization) as well as their interaction. Block (nested in treatment), block × year and family were random effects. The family effect was nested in years because we were interested in estimating the genetic variance within population each year of collection. The interaction between family and treatment was included in the family effect as a vectorial random effect. The complete model is summarized in equation [1], where *Y* denotes the flowering time, *μ* the average flowering time over the whole sample and *ϵ* the residuals:

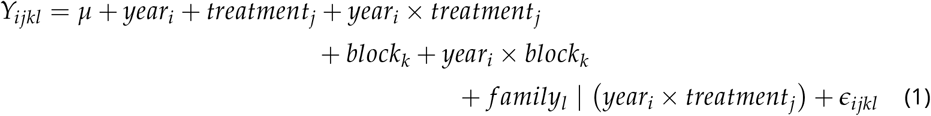

This maximal model was simplified, using likelihood ratio tests (LRT) to compare the models. In addition, we tested for a significant change in genetic variance between 1987 and 2009 using a LRT between the model [1] and a model where family is not nested into year. Standard errors for variance components were estimated using jackknife resampling. We used the variance components estimated for the random effects to calculate broad-sense heritability as 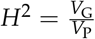, where *V*_G_ is the genetic variance as estimated by the family effect and *V*_P_ is the total phenotypic variance, including block, family and residual variance. Standard errors for *H*^2^ were estimated through jackknife resampling on families (Sokal and Rohlf, 1995).

### Temporal changes in sensitivity to vernalization

Selection on a trait in an environment can shift both the mean and the plasticity of that trait. Here, we considered the sensitivity to vernalization cues, measured as the slope of the regression line between individual values and the environmental value (estimated as the average phenotype, 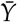) (Falconer and Mackay, 1996), for each individual *i*:

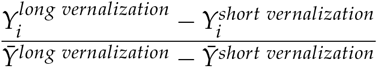

For each family, the five individuals in each treatment were paired according to their position in the greenhouse (block 1 with block 5, etc). This coefficient assumes that reaction norms are linear (Gavrilets and Scheiner, 1993; Scheiner, 1993) and this approximation is expected to work well (Chevin et al., 2013). We used a linear mixed model, with sampling year (1987 or 2009) as a fixed effect, a random block effect and its interaction with year, and a family effect (genetic effect) nested into year. As for flowering time, we estimated the broad sense heritability of the vernalization sensitivity.

A genetic correlation between flowering time and sensitivity to vernalization would affect the response to selection in the context of climate change. We therefore used a bivariate model with the sensitivity to vernalization and the flowering time measured in the short vernalization treatment as two dependent variables to estimate their genetic covariance with a random family effect, including block as a random effect, using AsReml (Gilmore et al., 2009). We ran an independent model for each sampling year. The significance of genetic covariances was tested by comparing the residual deviance of the final model with that of a model with a fixed covariance of zero in a log-likelihood ratio test.

### Selection gradient for flowering date: genetic covariance analysis

In the absence of selection for the trait considered, its observed variation is expected to be independent from fitness. We tested this by measuring the selection gradient, i.e. the statistical relationship between a trait and the fitness. Selection gradients were established for each year (and per treatment) following the Robertson-Price identity that states that Δ*Z*, the expected evolutionary change in the mean phenotypic trait *z* per generation is equal to Θ_*a*_(*z*, *w*), the additive genetic covariance of the trait *z* and the relative fitness *w* (Price, 1970; Robertson, 1966):

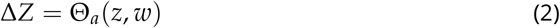

Here, we estimated the broad sense genetic covariance Θ_*g*_. Assuming that the dominance variance is negligible due to the very high levels of homozygosity in selfing populations (Holland et al., 2010), genetic covariance should be a good approximation of the additive genetic covariance (we neglect maternal genetic effects here). As a preliminary step, we checked whether our proxy for fitness, the relative seed production, had significant genetic variance. The relative seed production was measured as the individual seed production standardized by the average seed production of individuals from the same year and treatment. A mixed model was used to analyse the relative seed production, including two random effects for block and family. Then, provided there was significant genetic variance for relative seed production in the population each year, we analysed it in a bivariate model with flowering time to estimate the genetic covariance with a random family effect, including block as a random effect, using AsReml (Gilmore et al., 2009). Again, the significance of genetic covariances was estimated by comparing the residual deviance of the final model with that of a model with a fixed covariance of zero in a log-likelihood ratio test.

### Genetic analyses

During the multiplication generation in the greenhouse (2011), 200 mg of leaves were sampled from each plant for DNA extraction, using DNeasy Plant Mini Kit (Qiagen). Twenty microsatellite loci were used for genotyping (see the details of amplification reactions and analyses of amplified products in Jullien et al., 2019; Siol, Bonnin, et al., 2007). Briefly, samples were prepared by adding 3 *μ*l of diluted PCR products to 16.5 *μ*l of ultrapure water and 0.5 *μ*l of the size marker AMM524. Amplified products were analyzed on an ABI prism 3130 Genetic Analyzer and genotype reading was performed using GeneMapper Software version 5.

### Single-locus analyses assuming independence among loci

As a preliminary step, the data was filtered to reduce the percentage of missing data (loci or individuals with >10% missing data were removed), and to discard monomorphic loci. After filtering, the dataset comprised 145 individuals (representing 145 families) and 16 loci (64 from the year 1987 and 81 from the year 2009). We measured the genetic diversity of the population each year using the allelic richness *N*_*a*−*rar*_ (Hurlbert, 1971) and the expected heterozygosity *H_e_*. In this predominantly selfing population, we expect a strong deviation from Hardy-Weinberg heterozygosity expectations. Thus, for each sampling year, we estimated the inbreeding fixation coefficient *F_IS_* and its confidence interval using 5,000 bootstraps over loci. Between year differences for *N*_*a*−*rar*_, *H_e_* and *F_IS_* across loci were tested using Wilcoxon signed-rank tests. Analyses were performed in R using the packages adegenet (Jombart, 2008)and hierfstat (Goudet, 2005) and the program ADZE for rarefaction analyses (Szpiech et al., 2008). The percentage of pairs of loci showing significant linkage disequilibrium (LD) was calculated using Genepop (Rousset, 2008) with a threshold of 0.05. Finally, we measured the temporal variance in allele frequencies using the *F_ST_* estimator by Weir and Cockerham (1984). To estimate the effective population size (*N_e_*, measured in number of diploid individuals) from the temporal variance of allele frequencies, we used *F_ST_* estimates to account for the correlation of alleles identity within individuals due to selfing (Navascués et al., 2020) and followed the method outlined in Frachon et al. (2017). We measured a confidence interval for *N_e_* using an approximate bootstrap method (DiCiccio and Efron, 1996) over loci.

### Analyses based on multilocus genotypes

We used the program RMES to estimate selfing rates from the distribution of multilocus heterozygosity (David et al., 2007). We tested for a difference in selfing rates between years using a likelihood ratio test between models where the selfing rate was constrained to be constant or not. For each sample (1987 and 2009), we examined the genetic structure by sorting out the number of multilocus genotypes (thereafter called MLG) and measuring their frequency and redundancy through time using GENETHAPLO (available on GitHub at https://github.com/laugay/GenetHaplo and described in Supplementary Material S1). GENETHAPLO takes into account the uncertainty of the assignment of a genotype to a MLG group due to missing data: in case of ambiguity, an individual is randomly assigned to one of the candidate MLG group with a probability proportional to the MLG group size. The approach also considers a genotyping error rate: if two individuals differ by less than the error rate, they are considered to belong to the same MLG. After an initial run with an error rate of zero, we checked the distribution of the distances between MLGs. We found an excess of small distances, which could indicate errors in genotype assignation (Arnaud-Haond and Belkhir, 2007). We corrected this by re-running the program with an error rate of 1/16 (= one mis-read locus). GENETHAPLO also searches for residual heterozygosity (defined as the proportion of heterozygous loci in the multilocus genotype) and evidence for recombination (S1). To identify putative recombination events between MLGs, it uses the genetic distances: a MLG is a recombinant candidate if the sum of its allele differences with two other MLGs (“parental MLGs”) equals the number of allele differences between these two parental MLGs.

If a MLG has a high fitness in a given environment, plants carrying this MLG will produce on average a larger progeny and the frequency of the MLG will rise in the following generations. We therefore propose to use the absolute change in frequency of the fully homozygous MLGs through time as an indicator of their “realised fitness”. As a preliminary step, we checked whether selection quantified in the greenhouse is likely to mirror the predominant selection between 1987 to 2009 using a linear model to verify whether the change in MLG frequencies covaries positively with and can be predicted by the seed production in the greenhouse. We then measured the selection gradient for flowering time as the slope of the regression of the change in frequency of the MLGs between 1987 and 2009 with the genetic value of flowering time (measured as the average flowering time for a given MLG in the short vernalization treatment). We compared this pattern with the predictions from the Robertson-Price selection gradient. The MLGs found in 2009 but absent in 1987 may have been undetected in 1987 due to low frequency, or may be recent migrants. Their change in frequency between 1987 and 2009 is thus necessarily positive and may not accurately reflect their realised fitness. We therefore reiterated these analyses using a dataset restricted to the MLGs present in 1987 only. For each of these models, we verified the normality of the residuals and estimated a confidence interval for the slope using profile likelihood confidence bounds.

In addition, we tested whether the change in frequencies of the MLGs reflects a response to selection or can be expected by drift alone. This was tested by simulating the effect of 22 generations of drift, using an extension to multiallelic data of the approach described in Frachon et al. (2017) and inspired by Goldringer and Bataillon (2004). Again, only the fully homozygous MLGs were kept for this analysis. We assumed complete selfing during the time interval so the whole genome behaves as a single super-locus. Details about the procedure used to simulate individual MLG frequency trajectories are provided in Supplementary Material S2. We simulated each generation of drift by drawing MLG counts from a multinomial distribution parameterized with the effective population size *N_e_* estimated from the temporal *F_ST_*, and the MLG frequencies in the previous generation. Note that this simulation assumes a generation time of one year and therefore neglects seed dormancy and that the presence of a seed bank would reduce the rate of genetic drift. After 22 simulated generations, we randomly sampled 75 individuals to estimate the frequencies of each MLG and measured the change in MLG frequencies across the 22 generations. This was repeated for 104 replicates in order to draw the distribution of the change in MLG frequency expected by drift alone. To account for the potentially large estimation variance for the *F_ST_* (as observed in the simulations performed in Supplementary Material S3), we examined the sensitivity of the analysis to the effective population size using a range of values (10 ≤ *N_e_* ≤ 500). Finally, we examined the simulated selection gradient as the relationship between the simulated changes in MLG frequencies through time and the genetic value of flowering time previously measured for each MLG, using a linear model. This provided us with a null distribution of the slopes of the regression between frequency change and flowering time, expected under drift alone. We then tested for the significance of the observed slope against the simulated distribution, by computing the proportion of the simulated slopes that were greater than the observed value.

### Regional analysis

Finally, we attempted to disentangle selection and drift by considering other populations located in the same geographical region as the focal population and therefore likely submitted to the same selective pressure due to climatic constraints (pattern test, as described in the Introduction). For this regional analysis, we used 16 populations of *M. truncatula* across Corsica that were sampled twice, once in the 80’s and again in the early 2000s (listed in Table S1). Samples consisted of around 100 pods collected along transects running across the populations. Seeds collected were stored in a cold room. In 2010, one pod randomly selected from each sample (80’s and 2000’s) was threshed and one plant per population per year was replicated through selfing in standardized greenhouse conditions. This greenhouse generation allowed suppressing potential maternal effects (as in the experiment with the Cape Corsica population) and resulted in 32 families (16 populations × 2 years) of full-sibs produced by selfing. In 2011, seeds from the 32 families were germinated following the same protocol as described earlier for the intra-population analysis, but with only one vernalization treatment at 5°C during seven days. Five plants for each family were then transferred to tables in the greenhouse according to a randomized block design (five blocks). We monitored the temperature and humidity and the flowering time for each plant.

Individual flowering times were converted in thermal time, in the same way as it was done for the intra-population analysis. Again, we used linear mixed models (lme4 package) to test for the effect of sampling year on flowering time. The model included a single fixed effect for the sampling year (1980s or 2000s). The block effect was included as a random effect, along with its interaction with sampling year. A random population effect was also included and replaced the “family” effect of Eq. 1 seen as there was only one family per year in this regional sample. The resulting model was written as:

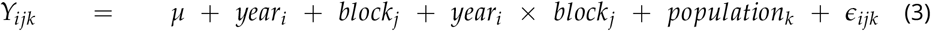

Again, this maximal model was simplified using likelihood ratio tests.

## Results

### Changes in flowering time

Visual inspection of the Q-Q plots indicated that the residuals from all the linear models we used were normally distributed. We found that flowering time differed significantly between years: plants sampled in 2009 flowered on average over two days earlier than plants sampled in 1987 (Table 1, Fig. 1). This effect remained significant when we analysed flowering time as a number of days rather than degree.days (results not shown). Longer vernalization also sped flowering up (treatment effect, Table 1). The block effect only explained a low proportion of variance (micro-environment) and the largest variance component was the family effect, for all combinations of years and treatments. The comparison of a model where family was nested in years only or in years × treatments showed that the family × treatment interaction was significant (*χ*^2^ = 66.1; df = 7; *p* = 9.10^−12^). It means that the reaction norms for the different genotypes were not parallel (Fig. 1), because the genotypes responded differently when exposed for a shorter period to cold temperatures. To account for this genotype × environment interaction, the heritability for flowering time was estimated in each vernalization treatment separately (four components of variance, Table 1). It varied between 0.53 and 0.77 (Table 2). The genetic variance for flowering time in the population remained the same in 1987 and 2009, as shown by a LRT between the full model (Eq. 1) and a model where family was not nested in year (*χ*^2^ = 6.65; df = 7; *p* = 0.47). We found no significant year effect on the sensitivity to vernalization (*χ*^2^ = 1.7; df = 1; *p* = 0.185). There was no significant difference in the family effect between years (interaction family × year not significant; LRT: *χ*^2^ = 1.2; df = 2; *p* = 0.552) but the family effect was highly significant (*χ*^2^ = 32.6; df = 1; *p* = 1.10^−8^, Table S2) and the heritability of the sensitivity to vernalization was 0.19 (+/− 0.04) (Table 2). Finally, the multivariate analysis highlighted a strong positive genetic correlation between flowering time (measured in the short vernalization treatment) and the sensitivity to vernalization (in 1987: 0.54 *p* = 0.008; in 2009: 0.60 *p* < 0.0001), which means that early flowering plants are less sensitive to vernalization cues. Using the flowering time measured in the long vernalization treatment, we observed the same pattern of correlation.

**Table 1.** Effect of sampling year and treatment on flowering time in the cape Corsica population, taking into account the family effect (genetic effect). Effect values on mean flowering time are given for fixed effects and variance components are given for random effects (with standard errors in brackets). The family effect was nested into year (1987 or 2009) and treatment (T1: short vernalization treatment; T2: long vernalization treatment), leading to four variance components. For each component, the degrees of freedom, likelihood ratio (*χ*^2^) and *p*-values are reported. None of the interactions considered in the complete model [1] were significant: between year and treatment (LRT *χ*^2^ = 1.8; df = 1; *p* = 0.178); between block and year (*χ*^2^= 0.0006; df = 1; *p* = 0.981).

| Tested effect on flowering time | Mean effect or variance component (SE) | df | $\chi^2$ | $p$ |
| --- | --- | --- | --- | --- |
| year | -28.76* | 1 | 7.3 | 0.007 |
| treatment | -162.84 | 1 | 42.2 | $8.10^{-11}$ |
| block | 92.34 (9.61) | 1 | 34.5 | $4.10^{-9}$ |
| family \| year × treatment | 1987-T1: 2807.90 (872.97) | 10 | 850.4 | $2.10^{-16}$ |
|  | 1987-T2: 1793.51 (500.25) |  |  |  |
|  | 2009-T1: 5449.80 (1200.16) |  |  |  |
|  | 2009-T2: 3557.01 (1408.88) |  |  |  |
| error | 1500 (38.73) | 1081 |  |  |

\* assuming an average daily temperature of 15°C over the time period considered, the difference of 28.76 degree.days corresponds to two days.

**Figure 1.**
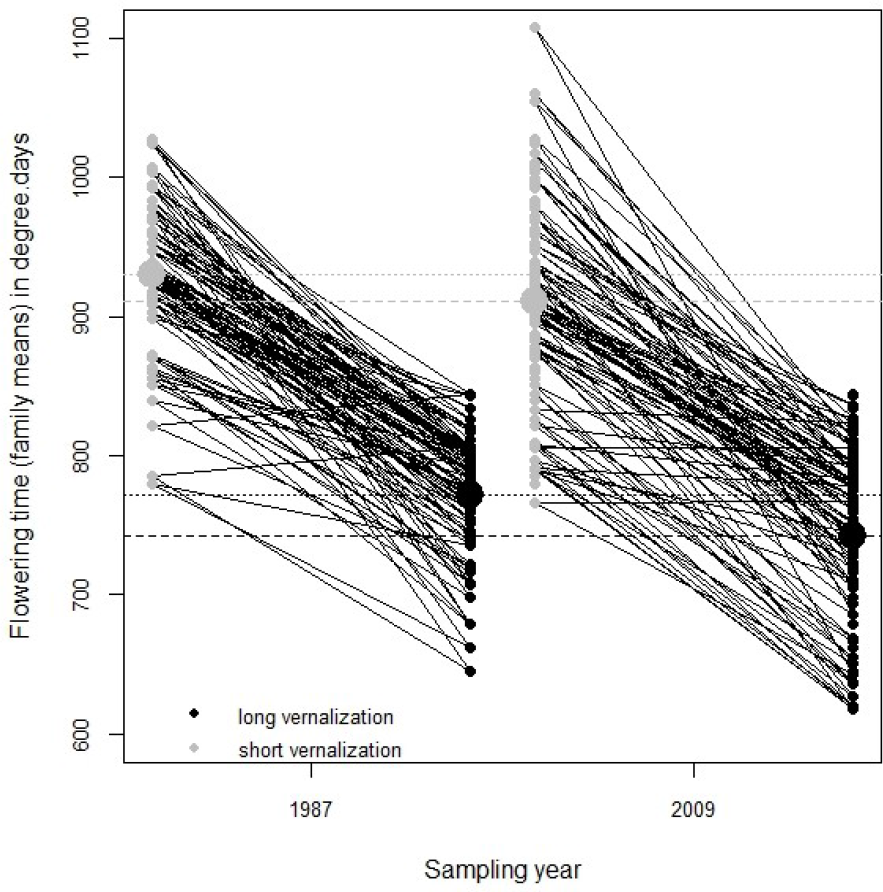
Average flowering time per family for the two sampling years and the two vernalization treatments. Short vernalization is in grey and long vernalization in black. The large dots and the horizontal lines stand for the average flowering date for each vernalization treatment, for the years 1987 (dotted lines) or 2009 (dashed lines). Black crossing lines indicate that the reaction norms differ between families, as expected if genotype × environment interactions are significant.

**Table 2.** Heritabilities (*H*^2^) and coefficients of genetic variance (*CV_g_*) for flowering time in each vernalization treatment (T1: short vernalization; T2: long vernalization) and each sampling year, for sensitivity to vernalization and for relative seed production.

| Trait | $H^2$ (SE) | | $CV_g$ | |
| --- | --- | --- | --- | --- |
|  | 1987 | 2009 | 1987 | 2009 |
| Flowering time | T1: 0.64 (0.06) | T1: 0.77 (0.04) | 5.70 | 8.11 |
|  | T2: 0.53 (0.07) | T2: 0.69 (0.07) | 5.49 | 8.03 |
| Sensitivity to vernalization | 0.19 (0.04) |  | 18.14 |  |
| Relative seed production | 0.34 (0.03) |  | 30.00 |  |

### Selection gradient for flowering date

The relative seed production showed significant genetic variance (family effect, Table S3, heritability of 0.34, Table 2), which enabled us to build multivariate models to examine selection gradients following Eq. 2. In 1987, we found a significant genetic covariance between flowering time and relative fitness: Θ_*a*_(*z*, *w*) = −20.5; LRT comparing this model with a model where the genetic covariance was constrained to be zero: *χ*^2^ = 60.2; df = 1; *p* = 8.10^−15^. The covariance remained significantly different from zero when we used the lines derived from the sampling in 2009: Θ_*a*_(*z*, *w*) = −18.5; LRT: *χ*^2^ = 12.4; df = 1; *p* = 6.10^−7^. A similar negative relationship was observed among lines derived from each of the two years, which means that the selection gradients predict an evolution towards early flowering under the environmental conditions of the greenhouse (Fig. 2).

**Figure 2.**
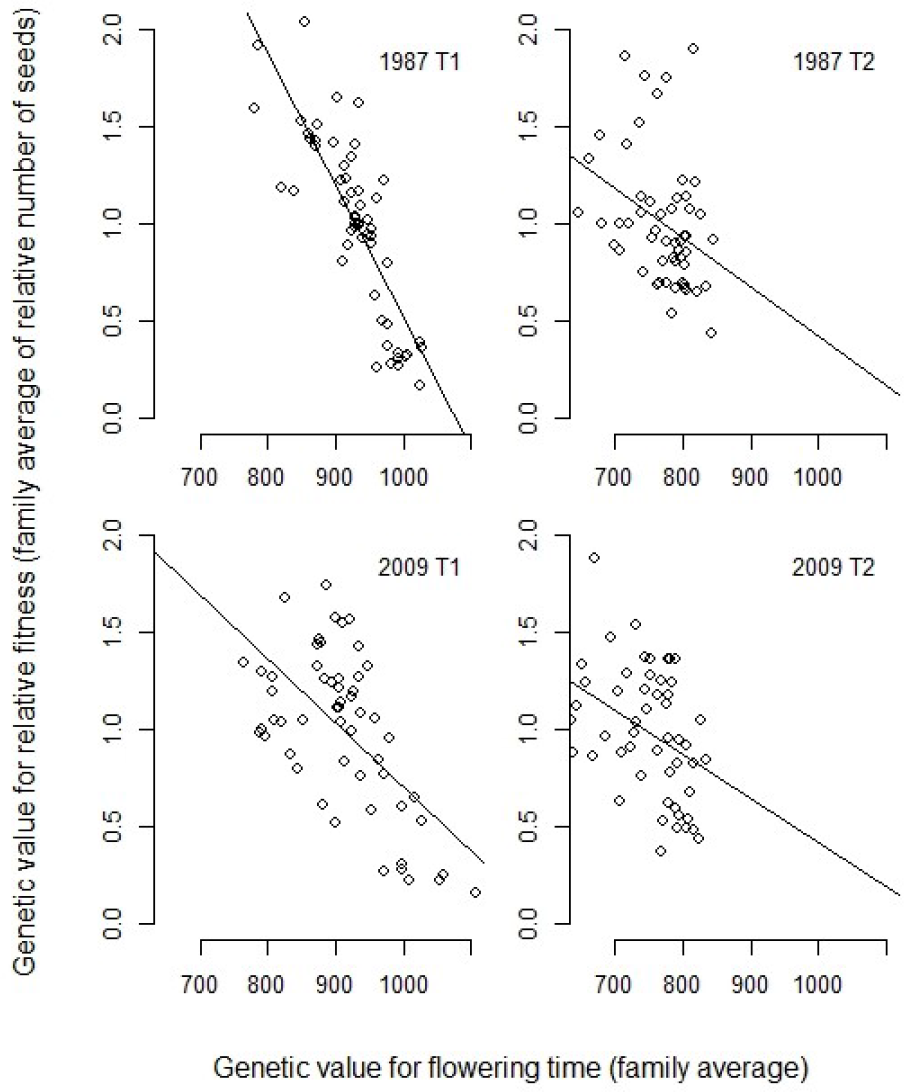
Selection gradients for flowering time. Established as the relationship between the genetic value for flowering time (family average, in degree.days) and the genetic value for relative fitness (family average of the relative number of seeds), for each sampling year and vernalization treatment. Lines stand for the linear regression.

### Changes in the genetic composition of the population

The analysis of microsatellite data highlighted high levels of genetic diversity for both sampling years, with an increase between 1987 and 2009 only significant for *H_e_* (Table S4). This suggests that the increased diversity between 1987 and 2009 reveals more balanced allele frequencies rather than an increase in the average number of alleles. The temporal differentiation measured using the 16 loci was high (*F_ST_* = 0.226; 95% confidence interval: 0.182 – 0.269), which translates into a particularly small effective size (*N_e_* = 19 diploid individuals; 95% confidence interval: 15-25). According to [Eq. 16] in Nordborg and Donnelly (1997), we predict that 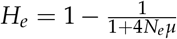, where *N_e_* is the effective size as estimated above. Using mutation rates for dinucleotide microsatellite loci measured in *Arabidopsis thaliana* (5.10^−5^ to 2.10^−3^) (Marriage et al., 2009), and assuming an isolated population at equilibrium, we expect that *H_e_* should lie between 0.004 and 0.134, which is nearly three times lower than the *H_e_* estimated here (Table S4). The observed heterozygosity was particularly low, resulting in large *F_IS_* estimates, as expected for a predominantly selfing species. The estimated selfing rate was about 94% in 1987 and rose to 98% in 2009 (statistically significant increase, Table S5). This high selfing rate results in extensive linkage disequilibrium between loci (nearly all pairs of loci are in linkage disequilibrium, Table S4), which makes the analysis of multilocus genotypes particularly relevant.

The analysis of MLG identified 60 different MLGs in this sample of 145 individuals. Out of the 60 MLGs, 48 were fully homozygous at the 16 loci and 12 MLGs displayed some level of heterozygosity (Fig. S1). We found no evidence for a link in terms of recombination or segregation between the heterozygous MLGs and any of the fully homozygous MLGs. These heterozygous MLGs were therefore excluded from the following analyses, leaving us with 48 MLGs (58 individuals in 1987 and 75 in 2009). The two predominant MLGs represented more than 50% of the population in 1987 and nearly 20% in 2009. These, as well as three other MLGs, were observed in both years (Fig. S2). The absolute changes in homozygous MLGs frequencies through time tended to covary positively with the total number of seeds produced by a plant in the greenhouse (Fig. 3A, regression only significant with the sample restricted to the MLGs present in 1987, *n* = 12 MLGs), which provides support to use it as a proxy to estimate the realised fitness. We therefore used the change in frequency of the 48 MLG (58 individuals in 1987 and 75 in 2009) to build selection gradients for flowering time. Again, we found a gradient with a negative slope (Fig. 3B), suggesting that the late flowering MLGs have a reduced realised fitness compared to earlier ones. This confirms the reduced fitness of late flowering genotypes observed in our greenhouse experiment (Fig. 2). Yet, the effect of flowering date on the realised fitness was small and only significant when the dataset was restricted to the MLGs present in 1987 and measured in the short vernalization treatment (*n* = 12; Fig. 3B). In addition, the negative slope was mostly supported by the decreasing frequency of the two late flowering MLGs that were prevalent in 1987. The simulation of 22 years of drift with an effective population size of 19 showed that the slope of the observed selection gradient did not deviate significantly from the distribution expected by drift alone (*p* = 0.182). Yet, again, when we restricted the dataset to the MLGs that were present in 1987, the observed selection gradient deviated significantly from the distribution expected by drift alone (*p* = 0.047), which suggests that the drift-alone hypothesis could be rejected.

**Figure 3.**
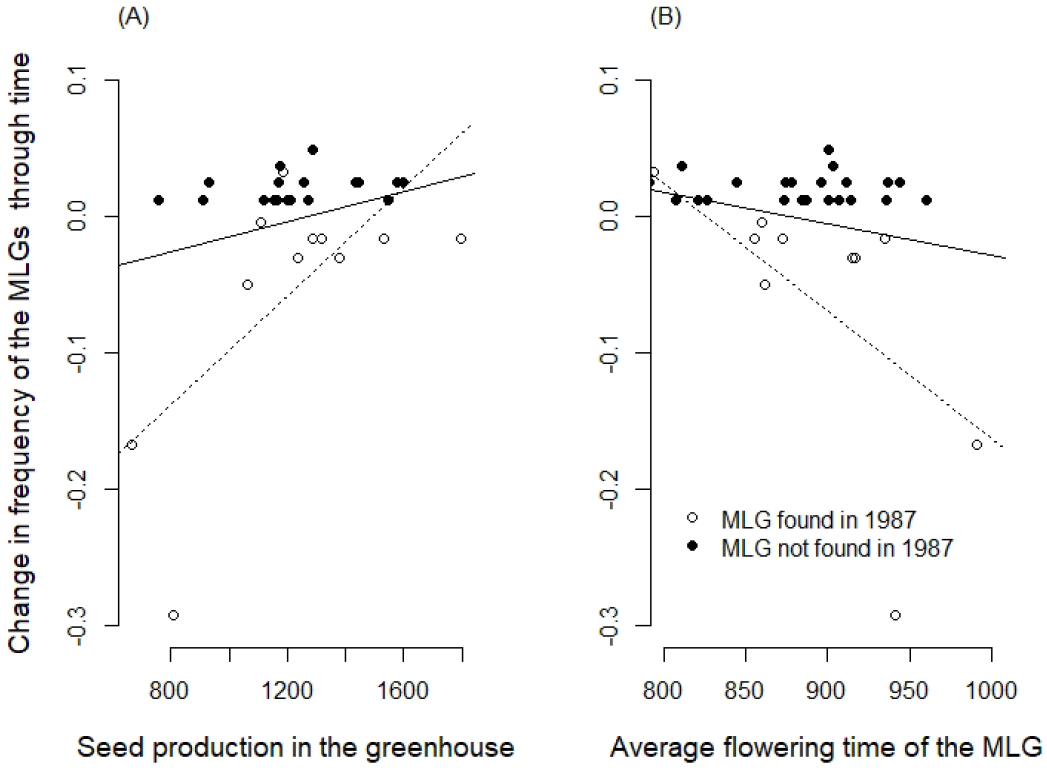
Analyses of the “realised fitness”, estimated as the absolute change in frequency of the MLGs through time. MLGs with residual heterozygosity were removed from this analysis. (A) Relationship with the average number of seeds produced by plants of a given MLG in the greenhouse. (B) Selection gradient for flowering time. Each point stands for the average flowering date for a given MLG. The black regression lines are estimated using all points (*n* = 48; A: slope = 5.10^−5^ points of frequency per seed *p* = 0.094; B: slope = −0.0002 95% confidence interval: −0.0006;0.0001 *p* = 0.179). This includes MLGs that were not observed in 1987 (black dots), for which the change in frequency is necessarily always positive. The dotted lines are the regression lines for the analysis restricted to the MLGs present in 1987 (white dots only; *n* = 12; A: slope = 0.0002 *p* = 0.024; B: slope = −0.0009 95% confidence interval: −0.0017;−0.0002 *p* = 0.038). Q-Q plots for the selection gradients are provided in Fig. S3.

Because selfing reduces the effective recombination, it reduces the number of independent loci. Measuring *F_ST_* from linked loci therefore amounts to measuring it from a lower number of markers, and it is known that *F_ST_* estimates based on a few loci suffer from a large sampling variance (Weir and Hill, 2002). Alternatively, we could have concatenated the genotypes at the different loci to compute a diploid version of the haplotype-based *F_ST_* (Mehta et al., 2019). Using the changes of frequencies for 48 homozygous MLGs, we estimated a temporal *F_ST_* of 0.075, which corresponds to an estimated effective size of 136. However, our simulations (Supplementary Material S3) show that these haplotype-based *F_ST_* estimates are strongly downwards biased, due to the dependency of *F_ST_* with allelic diversity (Alcala and Rosenberg, 2017; Edge and Rosenberg, 2014; Jakobsson et al., 2013) and could therefore overestimate the effective population size. Instead of using this unreliable estimate of 136, we assessed the sensitivity of our neutrality test for MLG frequency changes to the effective population size estimates, using a range of values (10 ≤ *N_e_* ≤ 500). We found that the observed selection gradient can no longer be explained by drift alone if the effective population size exceeds 150 (or even 10 if we consider only the MLGs present in 1987, Fig. 4).

**Figure 4.**
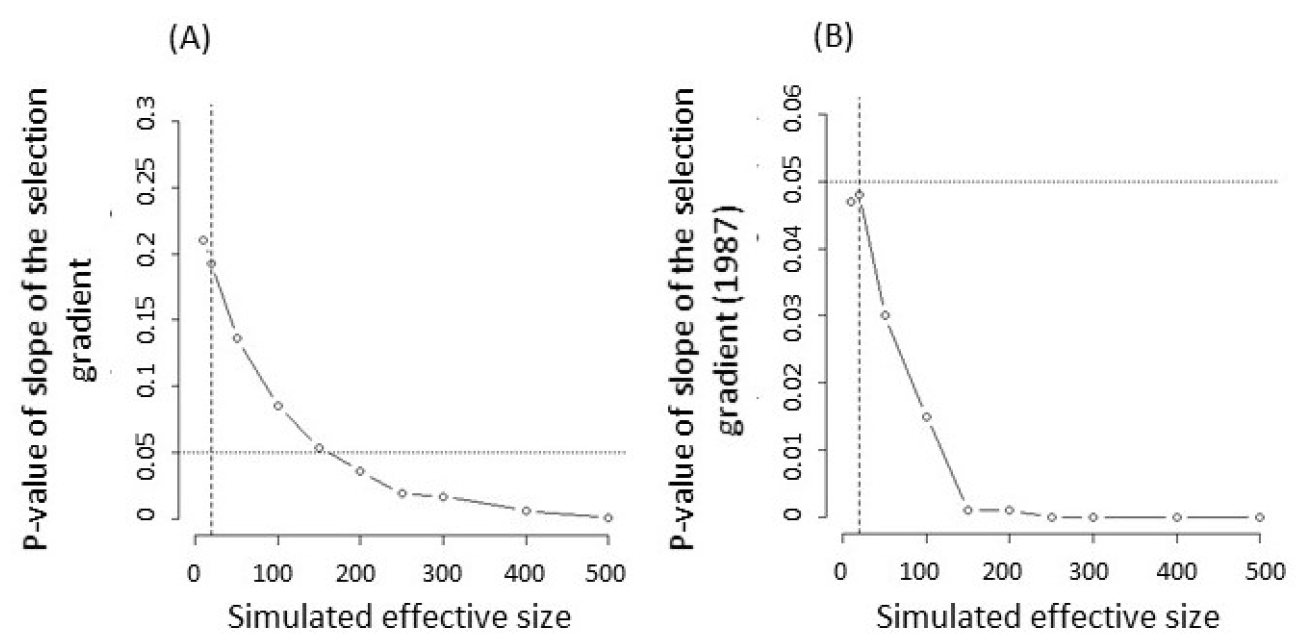
Test of selection for increasing values of *N_e_*. *P*-value, defined as the proportion of simulated datasets where the slope of the selection gradient is steeper than the observed slope, for the simulations of drift-alone (A) considering all the homozygous MLGs (*n* = 48) or (B) considering only the MLGs that were already present in 1987 (*n* = 12). The dotted line indicates the 0.05 threshold value for significance. The vertical dashed line is the effective size estimated using the temporal *F_ST_* and considering the 16 microsatellite loci as independent (*N_e_* = 19; *p* = 0.182 with *n* = 48 (A); *p* = 0.047 with *n* = 12 (B)).

### Changes in flowering time at the regional level

At the regional level (Eq. 3), we found no effect of the interaction between block and sampling year (LRT *χ*^2^ = 0; df = 1; *p* = 1). All other effects were significant (Table 3): the random block effect only explained 5% of the total variance whereas the population effect accounted for 34% of variance. The significant year effect showed that the material we collected in 2005 or 2009 in Corsica flowered about five days earlier (78 degree.days, Table 3) compared to the one we collected between 1987 and 1990.

**Table 3.**
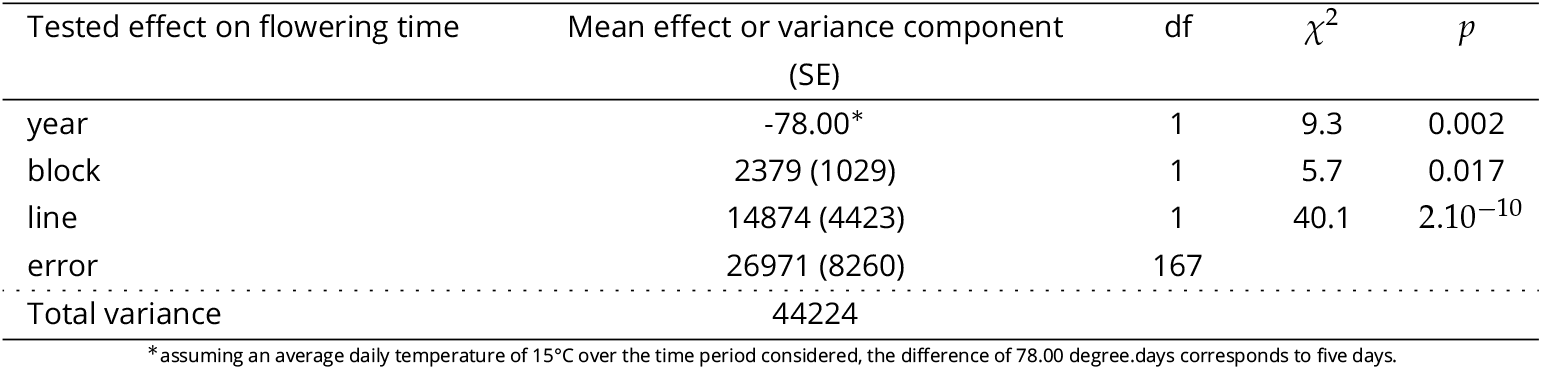
Effect of sampling year on flowering time at the regional scale, taking into account the effect of the population of origin of each line. The effect on the mean flowering time is given for the fixed year effect and variance components are given for random effects (with standard errors in brackets). For each component, the degrees of freedom, likelihood ratio (*χ*^2^) and *p*-values are reported.

## Discussion

Pairing up a resurrection study with population genetic analyses proved highly insightful to understand how flowering time changed through time in *M. truncatula* and to get insights into the mechanisms involved. Growing plants collected in the Cape Corsica population 22 generations apart in a common garden experiment provided evidence for a diminution of flowering times by about two days (i.e. a reduction between 2 and 4% in flowering time). This study also highlighted the peculiar genetic structure of this highly selfing population, where some multilocus genotypes are persistent through time. This enabled us to measure the fitness of a genotype as its frequency change through time and to establish a multilocus selection gradient. We used this multilocus fitness measure as well as a fitness measure based on individual seed production in the greenhouse to estimate the selection gradient for flowering time. Both gradients predict evolution towards earlier flowering but only the selection gradient using seed production in the greenhouse as a proxy for fitness was significant. It should be kept in mind that the selection gradient could change if the plants were growing in their natural environment, due to potential Genotype x Environment interactions. Simulating evolution across 22 generations showed that the observed change in flowering time can be caused by drift alone, provided the effective size of the population is lower than 150. These analyses suffer from the difficulty to estimate the effective size in a highly selfing population, where effective recombination is severely reduced.

### Can we use effective population size estimates to test whether the genetic change is caused by selection or drift in a predominantly selfing population?

As pointed out in the Introduction, simulating drift is one of the methods to test whether selection has occurred, but it requires knowledge about the effective population size. Using changes in allele frequencies between 1987 and 2009 in a natural population, we estimated a temporal *F_ST_* of 22.6%, which corresponds to an effective size of 19 (95% confidence interval: 15-25). This estimate is several orders of magnitude lower than the census population size (> 2,000 individuals) and lower than expected given the observed levels of diversity (Nordborg and Donnelly, 1997). Similarly low effective population sizes have been estimated previously in other *M. truncatula* populations, based on the temporal variance in allele frequencies (Siol, Bonnin, et al., 2007), and attributed to the high selfing rate of this species. Yet, the observed levels of polymorphism are often incompatible with such drastically low effective sizes (see Fig. 3c in Hereford, 2009; Jullien et al., 2019). *N_e_* estimates are likely biased and/or imprecise, because some of the assumptions underlying the temporal method are violated, e.g.: isolation of the populations under scrutiny, absence of selection, independence of marker loci (Jullien et al., 2019). For example, the quick change in allele frequency caused by a migration event will be misinterpreted as strong drift because temporal methods estimate *N_e_* using the pace at which allele frequency changes and therefore underestimate it (Wang and Whitlock, 2003). In addition, strong selfing affects the precision of temporal *F_ST_* estimates because the number of independent loci is reduced (Supplementary Material S3). In our focal population, the whole genome behaves practically as a single locus, which limits the precision of our effective size estimates. Unfortunately, we show in Supplementary Material S3 that inferring effective size from the variation of MLG frequencies (i.e., considering a single, multiallelic super-locus) is unlikely to improve the quality of our estimates.

Finally, if selection occurs in a non-random mating population, it will exacerbate the Hill-Robertson effect and further reduce the effective size (Comeron et al., 2007). Indeed, selection will create heritable variance in fitness among individuals, thereby locally reducing *N_e_* (Barton, 1995; D Charlesworth and Willis, 2009; Robertson, 1961). In predominantly selfing species, due to drastically reduced effective recombination (Nordborg, 2000), selection will extend the reduction in diversity caused by the selective sweep to a larger proportion of the genome compared to a random mating population (Caballero and Santiago, 1995; Kamran-Disfani and Agrawal, 2014). With selection, the effective size estimated using the temporal variance in allele frequencies can therefore not be considered as a “neutral” effective size but rather reflects the combined effects of inbreeding and selection (Le Rouzic et al., 2015). Overall, due to the reduced effective recombination and potential migration, predominantly selfing populations can strongly deviate from the assumptions of the temporal method to estimate effective size and such estimates should be treated with caution (See Fig. 3 in Jullien et al., 2019).

If highly selfing organisms strongly deviate from the general assumptions of population genetics models, a major benefit, however, is that the temporal survey of MLGs provides a highly integrative measure of fitness, which is analogous to measures of genotype-specific growth rates in asexual organisms. Our results show that changes in frequencies of MLGs through time are positively correlated to the fitness measured as the seed production in the greenhouse (Fig. 3A). This relationship is not significant if we consider all the MLGs found in 2009, but this is not surprising considering the potentially strong environmental variance in the field and the approximation due to the possibility that a MLG that was absent in 1987 appeared within the 22 years time period. A larger sample size in 1987 or additional temporal samples could help improve this analysis. Despite these imprecision, such integrative estimates of fitness are highly valuable because of the difficulty to obtain lifetime measures of fitness in the field (Shaw et al., 2008), which are generally hindered by pervasive trade-offs between life history traits such as reproduction and survival (Ågren et al., 2013; Anderson et al., 2014).

### What selective pressure could have led to this genetic change in flowering time? Insights from ecophysiology

The evidence that the change in phenology observed in this population across 22 generations is the result of selection as opposed to drift remains equivocal. A further step towards evaluating whether selection is responsible for the genetic change observed is to characterize the potential selective pressure involved. Phenological changes associated to climate change have been reported in a large number of plants (Amano et al., 2010; Cleland et al., 2007; Parmesan and Yohe, 2003; Root et al., 2003). In this context, ecophysiological models of phenology are insightful to understand how climate change can affect traits such as flowering time (Chuine, 2000; Oddou-Muratorio and Davi, 2014). The phenological response to climate change is complex, because the promoting effect of increased temperatures opposes the influence of reduced vernalization (Wilczek et al., 2010). Ecophysiological models generally predict a plastic shift towards earlier flowering times, as long as vernalization is sufficient during winter (Morin et al., 2009). In agreement with these predictions, a meta-analysis exploring the phenological response to climate change in plant populations showed that phenotypic changes are mostly plastic, while evidence for genetic adaptation remains relatively scarce (Merilä and Hendry, 2014, and other references of Evolutionary Applications special issue, January 2014). However, a large part of the intraspecific variation observed in phenology is genetic (Hendry and Day, 2005) and the architecture of the network underlying flowering time variation is well described in some species such as *Arabidopsis thaliana* (Sasaki et al., 2018; Wilczek et al., 2010). How climate change will affect the genetic values of phenological traits remains uncertain. In a first hypothesis, we may assume that the phenotypic optimum for flowering time is not affected by climate change. We therefore expect a genetic change occurring in the opposite direction than that of the plastic response (Fig. 5A). This hypothesis resembles counter-gradient variation, which occurs when the genetic influence on a trait along a gradient opposes the environmental influence, resulting in reduced phenotypic variation across the gradient (Levins, 1969). Counter-gradients are widespread along geographical gradients, as shown by the meta-analysis by Conover et al. (2009), who found evidence for counter-gradient in 60 species and for co-gradients in 11 species. Therefore, assuming that the same mechanism observed across spatial gradients could occur in temporal gradients, we would expect the genetic response of flowering time to counter-balance the plastic response to climate change. This could be achieved for example with a genetic change increasing the base temperature *T_b_* (temperature below which the development is supposed to be nil).

**Figure 5.**
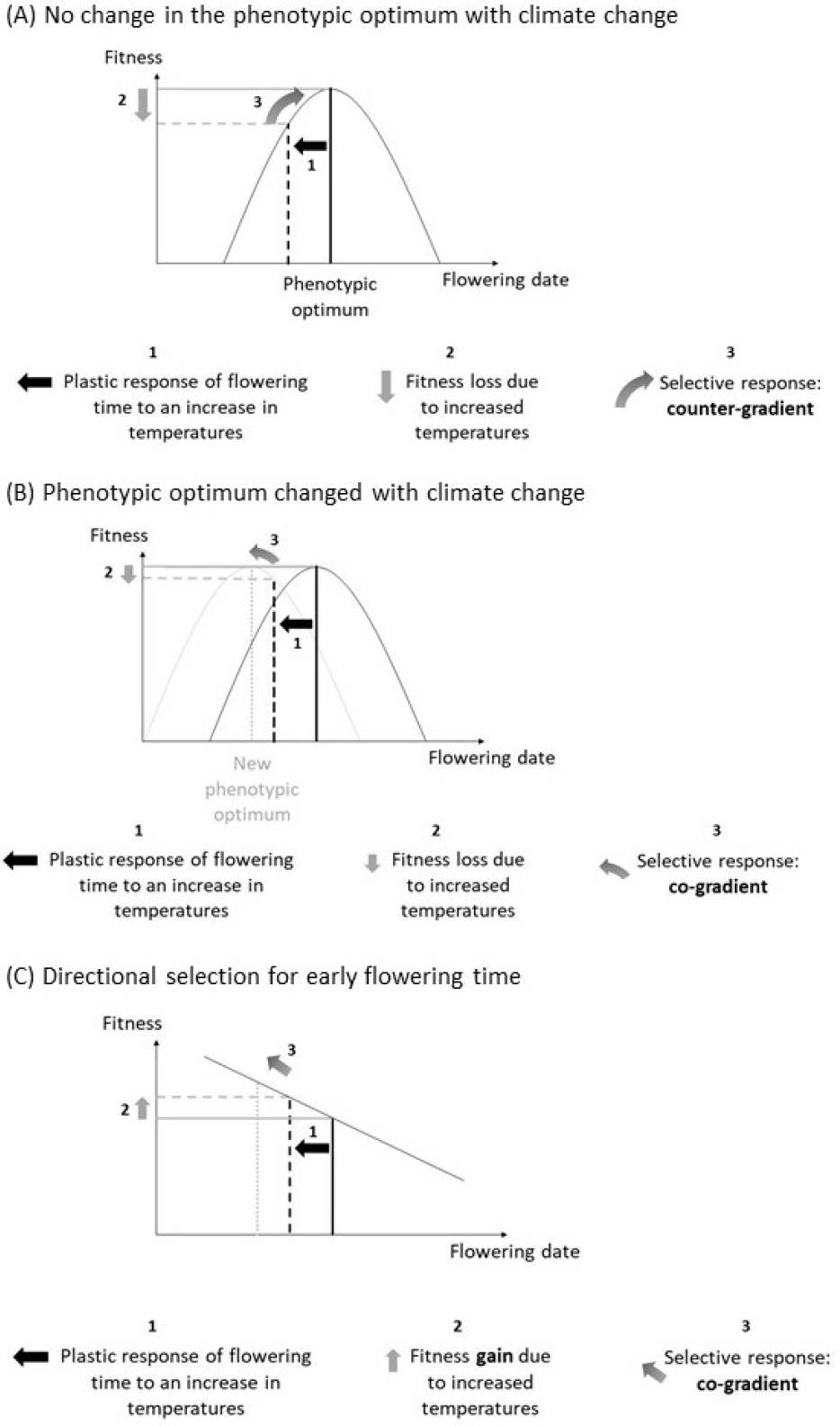
Hypotheses for the expected selective pressure on flowering time under climate change. (A) Selective response expected under the hypothesis that the phenotypic optimum for flowering date remains the same. The selective response is expected in the opposite direction compared to the plastic response to increased temperatures. This corresponds to the counter-gradient hypothesis. (B) Selective response expected under the hypothesis that the phenotypic optimum for flowering date is displaced with climate change and that it becomes advantageous to flower earlier. The selective response is expected in the same direction as the plastic response to increased temperatures. This corresponds to the co-gradient hypothesis. (C) Selective response expected under the hypothesis that flowering time is under directional selection.

Yet, our temporal survey rejects the counter-gradient hypothesis, both at the population and at the regional scale. Instead, we found evidence for a genetic change towards earlier flowering, in the same direction as the plastic response to the environmental change (here a rise in temperatures). Such a co-gradient is expected if climate change has shifted the phenotypic optimum towards earlier flowering dates (Fig. 5B). Several hypotheses could explain such a shift and the resulting co-gradient. First, in a plant with undetermined flowering such as *M. truncatula*, reduced frost risk early in the season should favour earlier flowering, because plants that manage to flower early in the season will carry on producing flowers until summer drought becomes limiting (end of May-June). We can therefore expect that the earliest a plant flowers, the highest its fitness. Second, climate change in the Mediterranean region also tends to reduce precipitations in spring and early summer (Goubanova and Li, 2007; Schröter et al., 2005), thereby shortening the reproductive period. Severe early summer drought could therefore create a strong selective pressure towards earlier flowering. Such a genetic shift in flowering time in response to extended drought have been reported before in the literature (Franks, Sim, et al., 2007). In terms of ecophysiology, it can be caused by lower requirements of degree.days, or a reduction of the base temperature *T_b_*.

Finally, although it is generally assumed that flowering date should be under stabilising selection in order to avoid frost or drought when flowering occurs respectively too early or too late, a recent meta-analysis found widespread evidence for frequent directional selection towards early flowering (Munguía-Rosas et al., 2011). Selection estimates considered in this meta-analysis largely ignore the effect of variation in number of flowers and plant size, which could bias the results. Yet, it remains that early flowering could have several advantages, among which an increased time for seed maturation in early reproducing plants and a longer period of growth for the progeny issued from seeds that germinate immediately (as reviewed by Elzinga et al., 2007; Kudo, 2006). Under this scenario of directional selection, we also expect a pattern of co-gradient, as observed in the data (Fig. 5C).

Besides the evidence for a genetic change in flowering date in *M. truncatula* in Corsica, we found no evidence for a change in the sensitivity to vernalization, despite genetic variance for this trait in the population (*H*^2^ = 0.19). In the literature, most studies have found at least some genetic variation for plasticity, but corresponding heritabilities were generally low (Scheiner, 1993). Our results also suggest that the sensitivity to vernalization is not independent from flowering date, because the intercept and the slope of the reaction norm to the vernalization treatment are genetically correlated (Gavrilets and Scheiner, 1993). Therefore, a lower number of chilling units received during winter (short vernalization treatment) results in higher heritability of flowering date. This correlation could favour the selective response of flowering date to climate warming because warmer winters will inflate the genetic variance of flowering date. Alternatively, if early flowering genotypes are selected for, or arrive in the population by migration, the evolution of the sensitivity to vernalization might be constrained by the positive genetic correlation with flowering time: early flowering genes tend to be associated with genes reducing the sensitivity to vernalization cues.

## Conclusions

Because it is difficult to rule out the effect of drift on the observed genetic change in phenology, our results do not entirely answer the question of the adaptive potential in selfing populations raised in the Introduction. Yet, several lines of evidence support the role of selection. First, the observed genetic change is in the direction expected for a response to raising temperatures and reduced rainfalls in the Mediterranean region. Second, the selection gradient measured in the greenhouse suggests that early flowering genotypes produce more seeds. The changes in MLG composition through time provide more equivocal results, but are also compatible with the hypothesis that MLGs with early flowering times had a better reproductive success than later flowering genotypes and replaced them, resulting in the observed genetic change in flowering time. Our simulations of the effect of drift are impacted by uncertainty in effective population size estimations, but the highest effective population size compatible with the observed change caused by drift alone remains relatively low (*N_e_* ≈ 150, Fig. 4A). Finally, the shift in flowering date observed in the Cape Corsica population was also detected at the regional scale, which suggests that the set of populations studied could be geographic replicates for this response to selection of flowering times in *M. truncatula* in Corsica. Ultimately, only a longer survey of this population combined with a pattern test (Sheets and Mitchell, 2001) could provide a definitive answer to the question of adaptation to climate change through a genetic change in flowering time in this predominantly selfing population. Finally, it is worth pointing out that, in contrast with the theoretical predictions presented in the Introduction, this population displays significant genetic variance for a quantitative trait such as flowering time. As suggested before for *M. truncatula* (Jullien et al., 2019), it is likely that other evolutionary mechanisms, such as migration, contribute to maintain the adaptive potential of populations in this predominantly selfing species.

## Data accessibility

Phenotypic data for the intra-population and inter-population experiments and results from the multilocus genetic structure along with the scripts used for the analyses are available on the INRA dataportal https://data.inrae.fr/dataset.xhtml?persistentId=doi:10.15454/ZY83BE. The program GENETHAPLO is available at https://github.com/laugay/GenetHaplo.

## Supplementary material

Supplementary material is available below and consists of the following. Table S1: List of the 17 populations sampled in Corsica (France). Table S2: Summary of the GLM on the sensitivity to vernalization. Table S3: Summary of the GLM on the relative seed production. Table S4: Genetic diversity at the 16 microsatellite loci. Table S5: Estimates of the selfing rate. Figure S1: Distribution of residual heterozygosity across MLGs for the two sampling years pooled. Figure S2: Distribution of MLGs in the population and through time. Figure S3: QQplot for the selection gradient in Figure 3B. Supplementary Material S1: Description of GENETHAPLO, a java program to analyse the genome-wide multilocus genetic structure of predominantly selfing or clonal populations. Supplementary Material S2: Details about the multiallelic method to simulate the effect of successive generations of drift. Supplementary Material S3: Comparison of the *F_ST_* estimation variance when considering the loci as independent or using the multilocus genotypes as alleles of a single locus. Figure S4 and S5: results of the simulations.

## Acknowledgements

The authors thank J.M. Prosperi for the collection of seeds as well as D. Tauzin and P. Noël for their help in running the greenhouse experiments. K. Loridon, C. Tollon, V. Lemaire and E. Figuet contributed to the production of the microsatellite data. This research was developed under the SelfAdapt project, funded by INRAE metaprogram “Adaptation of Agriculture and Forests to Climate Change” (ACCAF). Additional funding was provided by the Agence Nationale de la Recherche [ANR SEAD-ANR-13-ADAP-0011]. We are grateful to Christoph Haag, Pierre Olivier Cheptou, Jon Agren and Stefan Laurent for their comments on this work. Version 4 of this preprint has been peer-reviewed and recommended by *Peer Community In Evolutionary Biology* (https://doi.org/10.24072/pci.evolbiol.100128)

## Author Contributions

Joëlle Ronfort and Laurène Gay conceived and designed the research. JD conducted the greenhouse experiment in 2012 during his master’s project. MN, RV, LG and JR developed the extension to multi-allelic data of the method to detect selection using temporal data. MJ and LG ran the genetic analyses of the CO3 population. VR developed GENETHAPLO, the program to analyse the genome-wide multilocus genetic structure of selfing or clonal populations. LG wrote the article with the help of all authors that critically reviewed and approved the text.

---

**Table S1.** List of the 16 populations sampled in Corsica (France) with geographic coordinates and sampling years

| Population | Latitude | Longitude | Altitude | Sampling years |
| --- | --- | --- | --- | --- |
| FRA20025 | 42.756332397 | 9.4508333206 | 60 | 1985-2005 |
| FRA20031 | 42.182167053 | 9.3766670227 | 130 | 1985-2005 |
| FRA20035 | 42.361167908 | 9.4963331223 | 280 | 1985-2009 |
| FRA20039 | 42.021499634 | 8.7348337173 | 410 | 1985-2005 |
| FRA20043* | 42.462001801 | 8.6848335266 | 40 | 1987-2009 |
| FRA20044 | 42.551166534 | 8.7391662598 | 250 | 1985-2005 |
| FRA20046 | 42.592441559 | 8.9075956345 | 240 | 1985-2005 |
| FRA20049 | 42.444000244 | 9.4596662521 | 120 | 1985-2005 |
| FRA20050 | 42.165782928 | 9.546872139 | 10 | 1985-2009 |
| FRA20051 | 42.199165344 | 9.4646663666 | 100 | 1987-2005 |
| FRA20056 | 41.41350174 | 9.1668329239 | 60 | 1985-2009 |
| FRA20058 | 41.405334473 | 9.1265001297 | 195 | 1985-2009 |
| FRA20069 | 42.591667175 | 8.9081668854 | 410 | 1987-2005 |
| FRA20087 | 42.901668549 | 9.470000267 | 15 | 1987-2005 |
| FRA20088 | 42.958332062 | 9.3950004578 | 160 | 1987-2005 |
| FRA20089† | 42.970500946 | 9.3668336868 | 380 | 1987-2009 |

**Table S2.** Effect of sampling year on sensitivity to vernalization, taking into account the family effect (genetic effect). For each effect, the variance component (with standard errors in brackets), the deviance, degrees of freedom, likelihood ratio (*χ*^2^) and *p*-values are reported.

| Tested effect on sensitivity to vernalization | Variance component (SE) | df | $\chi^2$ | $p$ |
| --- | --- | --- | --- | --- |
| block | 0.01 (0.003) | 1 | 21.0 | $5.10^{-6}$ |
| family | 0.03 (0.008) | 1 | 32.6 | $1.10^{-8}$ |
| error | 0.13 (0.012) | 542 |  |  |

**Table S3.** Analysis of the family effect (genetic effect) on relative seed production (seed production standardized by year and treatment), taking into account the block effect. For each random effect, variance components (with standard deviations in brackets), degrees of freedom, likelihood ratio (*χ*^2^) and *p*-values are reported.

| Tested effect on relative seed production | Variance component (SD) | df | $\chi^2$ | $p$ |
| --- | --- | --- | --- | --- |
| block | 0.024 (0.15) | 1 | 116 | $< 2.10^{-16}$ |
| family | 0.090 (0.30) | 1 | 291 | $< 2.10^{-16}$ |
| error | 0.153 (0.39) | 1094 |  |  |

**Table S4.** Genetic diversity at the 16 microsatellite loci for the Cape Corsica population in 1987 and 2009. *n* stands for the sample size, *N_a_* and *N_arar_* are the average number of alleles per locus and the allelic richness (after correction using a rarefaction method), *H_e_* is the expected heterozygosity, *F_IS_* is the heterozygote deficiency (both with 95% confidence interval in brackets, estimated by bootstrapping the individuals) and *LD* is the percentage of loci under significant linkage disequilibrium (for a type I error fixed at 5% when rejecting the equilibrium hypothesis). Multilocus diversity is described by *n_MLG_*, the number of multilocus MLGs and 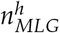, the number of fully homozygous MLGs. Wilcoxon signed rank tests were performed across loci for *N*_*a*−*rar*_, *H_e_* and *F_IS_* to test for a significant difference between the two years and the *p*-values are given.

| Sampling year | $n$ | $N_a$ | $N_{arar}$ | $H_e$ (CI95) | $F_{IS}$ (CI95) | $LD$ | $n_{MLG}$ | $n_{MLG}^h$ |
| --- | --- | --- | --- | --- | --- | --- | --- | --- |
| 1987 | 64 | 3.6 | 3.4 | 0.351<br>(0.252-0.424) | 0.942<br>(0.913-0.966) | 89% | 18 | 12 |
| 2009 | 81 | 3.9 | 3.7 | 0.623<br>(0.599-0.627) | 0.967<br>(0.957-0.976) | 96% | 47 | 41 |
| Total | 145 |  |  |  |  |  | 60 | 48 |
| $p$-value | | | 0.211 | 6.10-5 | 0.090 | | | |

**Table S5.**
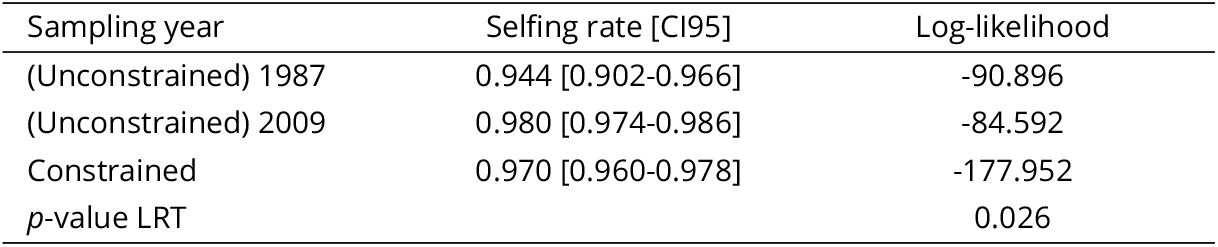
Estimates of the selfing rate in the Cape Corsica population for each sampling year obtained using the program RMES by maximizing the log-likelihood of the whole multilocus heterozygosity structure of the sample. The 95% confidence intervals and the log-likelihood are given for the two unconstrained models and the constrained model, along with the *p*-value of the likelihood ratio test comparing constrained and unconstrained models.

| Sampling year | Selfing rate [CI95] | Log-likelihood |
| --- | --- | --- |
| (Unconstrained) 1987 | 0.944 [0.902-0.966] | -90.896 |
| (Unconstrained) 2009 | 0.980 [0.974-0.986] | -84.592 |
| Constrained | 0.970 [0.960-0.978] | -177.952 |
| $p$-value LRT | | 0.026 |

**Figure S1.**
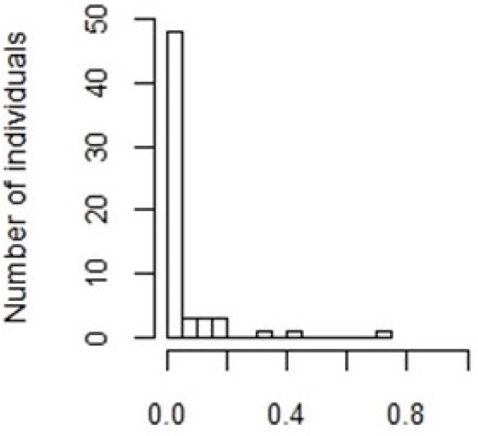
Distribution of residual heterozygosity across MLGs for the two sampling years pooled. Residual heterozygosity is defined here as the proportion of heterozygous loci in the multilocus genotype (over 16 loci) of each individual.

**Figure S2.**
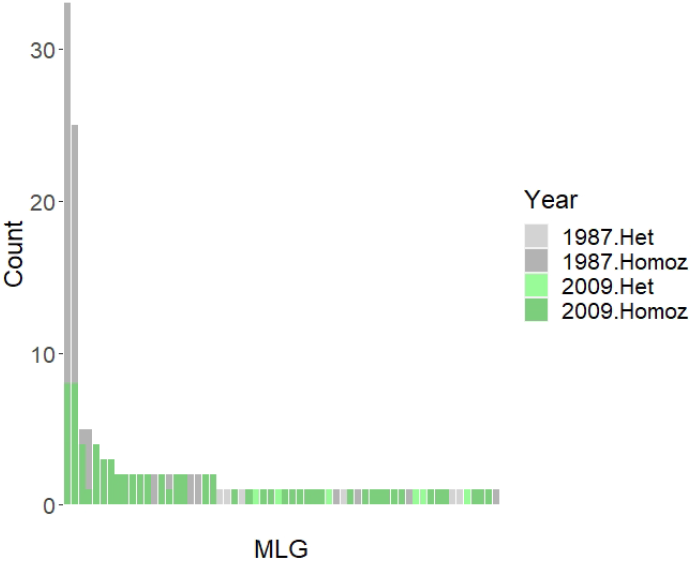
Distribution of MLGs in the population and through time. The four most frequent MLGs are shared between years. MLGs with residual heterozygocity are shown in light grey (for the year 1987) and light green (for the year 2009).

**Figure S3.**
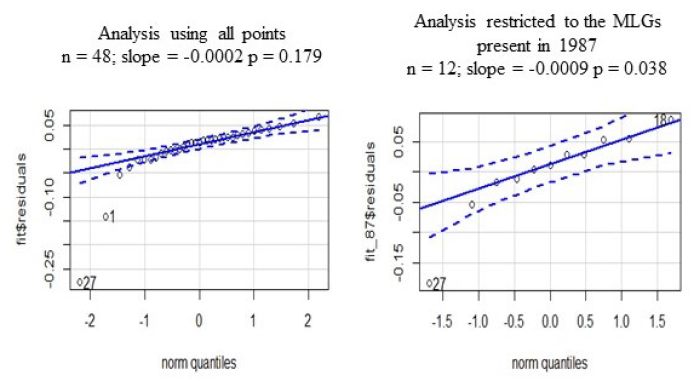
Q-Q plots for the selection gradients shown on Figure 3B.

## S1 GENETHAPLO: a java program to analyse the genome-wide multilocus genetic structure of predominantly selfing or clonal populations

Multilocus genotypes provide valuable information about mating systems (Jullien *et al.* 2019). Four software packages were previously developed to identify individuals originating from clonal reproduction using their multilocus genotype: MLGSIM (Stenberg *et al.* 2003); GENOTYPE and GENODIVE (Meirmans and Van Tienderen 2004), GENECLONE (Arnaud-Haond and Belkhir 2007) and poppr (Kamvar *et al.* 2014). Yet, none of these programs is specifically designed to identify individuals reproducing by selfing, in particular to detect repeated multilocus genotypes within a population and through time (or space) and recognize potential recombinants, formed by rare outcrossing events. GENETHAPLO is a program written in Java with four modules:

1. A module to convert the format of a dataset
2. A module to filter the dataset
3. A module to analyse the genetic diversity
4. A module to analyse the multilocus genetic structure

### Formatting the data file

The first line of the data file is a header line describing the content of each column, i.e. the name of the population, of the sub-population, of the individual and of each locus. Each following line provides the genotype of an individual at the specified loci. The individuals should be sorted so that populations and sub-populations are grouped together in consecutive lines.

Example:

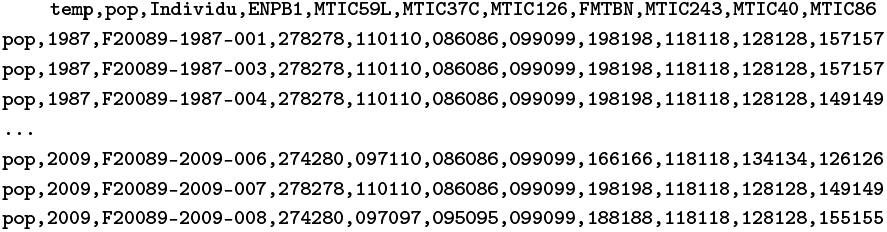

#### Module 1: format conversion

This module takes a dataset in the read2snp (Uricaru *et al.* 2014) format and converts it to a format suitable for GENETHAPLO, as detailed above.

#### Module 2: data filter

This module allows to filter out the loci, and individuals, having a percentage of missing data exceeding a specific threshold (given by the user). The two output files are i) a reduced dataset and ii) the list of the loci and individuals that have been removed. The percentage of missing data before and after filtering is also provided.

#### Module 3: genetic diversity

This module computes the key descriptors of genetic diversity classically used in population genetics studies. A first table summarizes the average number of individuals, alleles, the expected and observed heterozygosity and the *F_IS_*. The selfing rate is also calculated from the *F_IS_* for each sub-population. The module also provides these descriptors of diversity per locus and a table of allele frequencies for each sub-population.

#### Module 4: multilocus genetic structure

This module comprises three steps:

1. Grouping individuals according to their multilocus genotypes (thereafter called MLG): This module is based on a graph algorithm, where each node is an individual and nodes are connected when the individuals share the same MLG. An error rate can be specified by the user to allow grouping MLGs that differ at less than a given proportion of loci. This avoids over-splitting the MLGs due to genotyping errors or recent mutations. The module also takes into account missing data that can generate uncertainties. In case of missing data, it is possible for an individual to have a genotype compatible with several MLG. In such a case, the individual is randomly assigned to one of the possible MLG groups based on a random draw where each MLG group has a probability of being chosen that is proportional to its size. The output files provide i) the list of all individuals with the MLG to which they are assigned ii) the list of all identified MLGs with their frequency in each sub-population, their residual heterozygosity, defined as the proportion of heterozygous loci out of the total number of loci without missing values, and the number of missing values in each MLG.
2. Estimating genetic distance between MLGs: The genetic distance between two MLGs is estimated as the number of alleles that differ between the two synthetic MLGs divided by the total number of alleles without missing data in these two MLGs. This module generates a distance matrix as well as a histogram depicting the pairwise distance distribution.
3. Identifying recombinant MLGs: This module uses the genetic distances to rapidly identify putative recombination events between MLGs. A MLG is a candidate recombinant between two other MLGs (thereafter called “parental MLGs”) if the sum of the allele differences between it and its two putative parents equals the number of allele differences between these two parental MLGs. Only the MLGs that are represented by at least two individuals can be considered as potential parents. The output file provides a list of potential families, with the details of pairwise genetic distances.

### Running the program

This java program can be launched from a command prompt, in the folder where the modules are stored, using the command `java -jar module.jar`, where `module.jar` should be replaced by the corresponding module name.

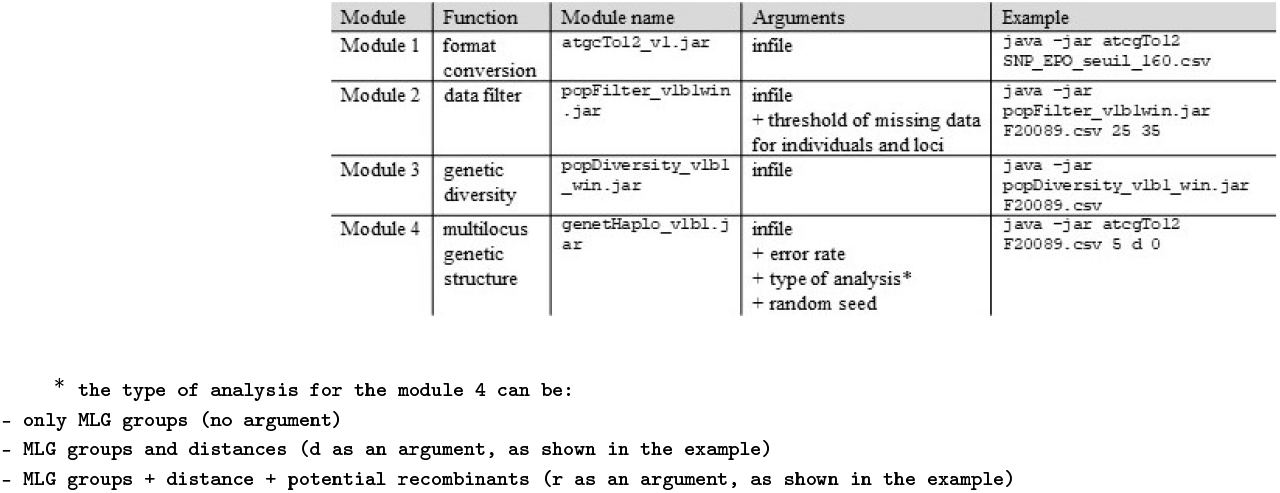

If no argument (infile or option) is added in the command, a short description of the script is displayed.

For example:

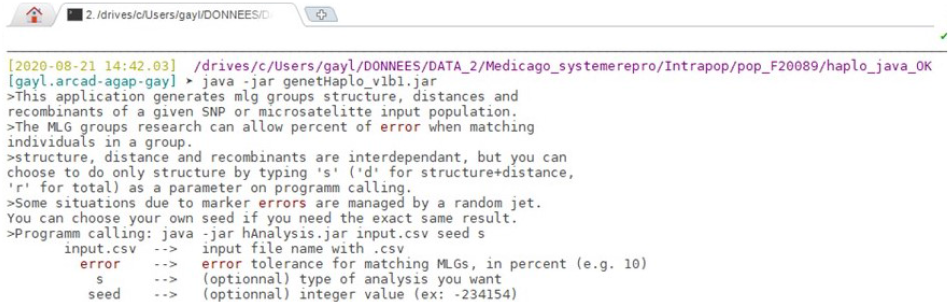

Example of output of the console:

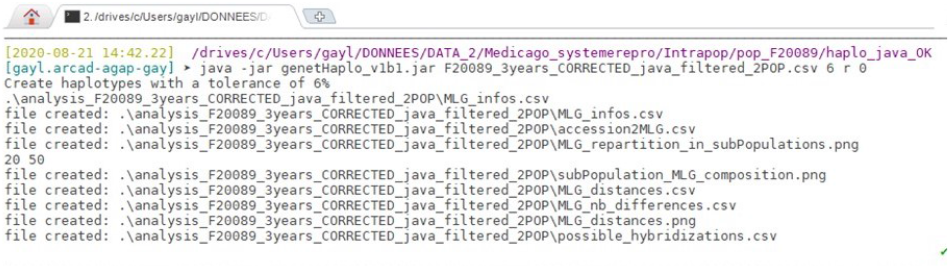

Example of output figures:

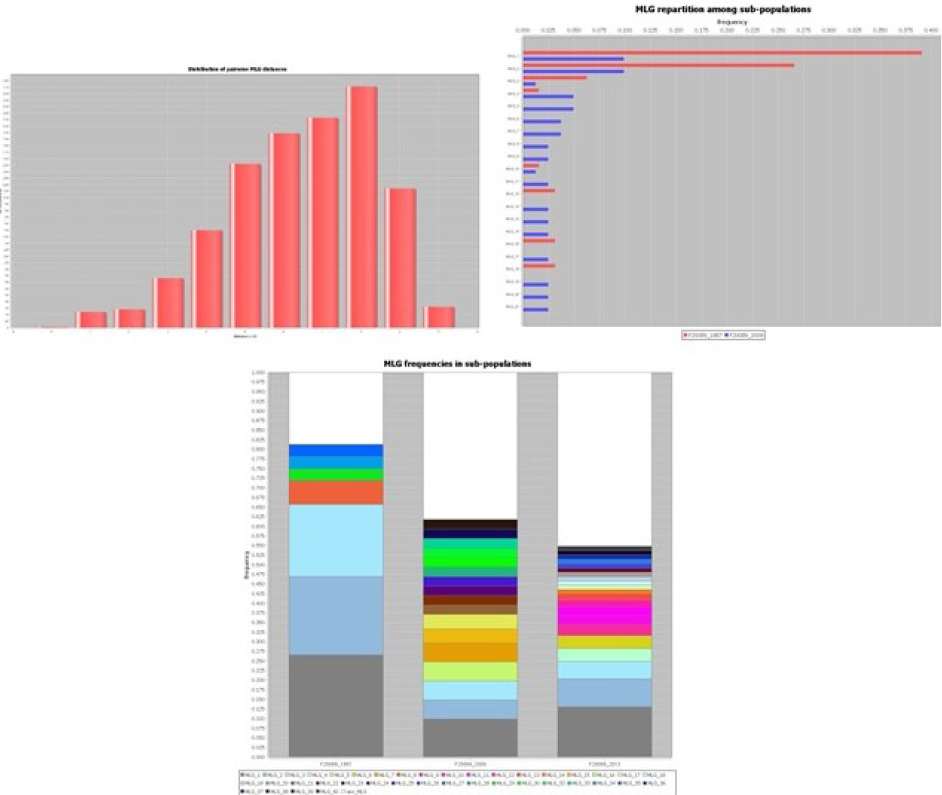

GENETHAPLO is a program freely available at https://github.com/laugay/GenetHaplo. Source codes are available from authors upon request.

## S2 Details about the multiallelic method to simulate the effect of successive generations of drift

We simulated the effect of 22 generations of drift, using an extension to multiallelic data of the approach described in Frachon *et al.* (2017) and inspired by Goldringer and Bataillon (2004). In order to account for the sampling variance in initial MLG frequencies, we simulated individual MLG frequency trajectories as follows: suppose that we observe a vector **y** of MLG counts, out of *n* total counts, in the 1987 sample. We assume that these observed counts are drawn from a multinomial distribution Mult(*n*, **x**) where **x** is the vector of (unknown) MLG frequencies in the 1987 population. Assuming a Dirichlet Dir(1) prior distribution for **x**, and using the Bayes inversion formula, the posterior distribution of **x** is distributed as Dir(**y** + 1). For each simulation, we therefore randomly draw the initial MLG frequencies in the 1987 sample ***π*_1987_**, from a Dir(**y** + 1) distribution. We then draw “pseudo-observed” MLG counts using a random draw from Mult(*n*, ***π*_1987_**).

## S3 Comparison of the *F_ST_* estimation variance when considering the loci as independent or using the multilocus genotypes as alleles of a single locus

Due to reduced effective recombination, the entire genome of a predominantly selfing population can behave as a giant supergene. This violates the hypothesis of independence between loci that is commonly assumed in population genetics, in particular for *F_ST_* estimation. One solution to this violation could be to take the linkage disequilibrium into account by concatenating all loci and considering the distinct multilocus genotype (thereafter MLG) as different alleles of a single (mega) locus. Here we use simulations to compare estimates of genetic differentiation measured using all loci considered independent or using MLGs as alleles of a single locus, and compare bias and estimation variance (MSE).

## Methods

We used the individual-based simulations of diploid hermaphroditic populations performed using SLiM 2.5 (Haller and Messer 2017) by Jullien *et al.* (2019). Briefly, we simulated the evolution of 20 independent loci (with a recombination rate of 0.5) in an isolated population with a given selfing rate and effective population size. Each simulation comprised two periods. A first period of 25 *N* generations (with *N* the demographic population size, measured as the number of diploid individuals) allowed the populations to reach the mutation-drift equilibrium. At this stage (time *t*_0_ = 0), 100 diploid individuals were randomly sampled. Twenty generations later (*t*_20_), a second sample of 100 individuals was drawn. This matches the sampling design performed on the focal population in Cape Corsica. Further details can be found in Jullien *et al.* (2019). We simulated 1,000 replicates for populations with a selfing rate ranging between 0.8 and 1 and a population size *N* of 250 individuals. For each simulated dataset, we assessed the relative temporal differentiation between the two samples using Weir and Cockerham’s (1984) estimator of *F_ST_*, as implemented in the R package hierfstat (Goudet 2005). We then grouped individuals with identical combinations of alleles (multilocus genotypes, MLG) using the program GENETHAPLO (Supplementary Material S1 above and as detailed in the main text, except for the error rate, which was set to zero). MLGs with residual heterozygosity were removed for the multilocus assessment of temporal differentiation. We considered each MLG as an allele of a single locus and computed the allele frequency for each temporal sample (*t*_0_, *t*_20_). We used the function haploDiv of the R package diversity (Keenan *et al.* 2013) to estimate the *F_ST_* on this haploid locus using Weir and Cockerham’s method (1984). We also reiterated this analysis without removing the MLGs with residual heterozygosity to assess the effect of this step on the bias and variance of *F_ST_* estimation.

For each simulated selfing rate, we calculated the expected *F_ST_* using the relationship established in Frachon *et al.* (2017): *F_ST_* = *τ*/(4*N_e_* + *τ*), where *N_e_* is the number of haploid individuals (or gene copy number) and *τ* the number of generations between the two temporal samples. Selfing reduces the number of independent gametes sampled for reproduction, so that the effective size is reduced to *N_e_* ~ *N*(2 − *σ*)/2 (Wright 1969, Pollak 1987) with *σ* the selfing rate and *N* the population size. As a result, 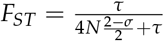.

We measured the bias as the difference between this reference *F_ST_* and the *F_ST_* we estimated assuming independent loci or the *F_ST_* we estimated using MLGs as alleles of a single locus. We measured the mean square error as the sum of the bias and the variance: 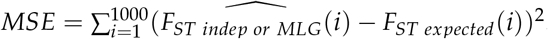.

## Results and discussion

When using MLGs as alleles of a single locus, the estimated *F_ST_* suffers from an increased negative bias compared to the method assuming that all loci are independent (Fig. S4). The bias decreases with increasing selfing rate but is always negative for the selfing rates we considered (>0.8), which will tend to overestimate the effective population size (Fig. S4). This bias is likely due to the dependence of *F_ST_* on the frequency of the most frequent allelic type (Jakobsson *et al.* 2013, Edge and Rosenberg 2014, Alcala and Rosenberg 2017): as the number of alleles increases, the frequency of the most frequent allele necessarily decreases, which sets an upper bound to the *F_ST_* estimates (Fig. 2 in Jakobsson *et al.* 2013). Removing MLGs with residual heterozygosity reduces the bias, because heterozygous MLGs are generally unique and therefore form new alleles of the single “MLG” locus. In addition, the precision of the *F_ST_* estimates using the MLG method is expected to decrease when the number of loci considered increases, because genetic diversity is influenced by haplotype length (Mehta *et al.* 2019). As already shown (Navascués *et al.* 2020), the variance of the *F_ST_* estimation assuming independent loci increases with high selfing rates (Fig. S5), due to the linkage disequilibrium that reduces the number of effective loci (Golding and Strobeck 1980, Nordborg 2000). Surprisingly, the MLG method seems to limit the estimation variance. This is most probably artificial, because the upper-bound on the *F_ST_* estimates also constrains its variance. Altogether, despite the high sampling variance due to the low number of effective loci available under strong selfing, our results suggest that it is preferable to assume that all loci are independent instead of using MLGs to estimate *F_ST_*.

**Figure S4.**
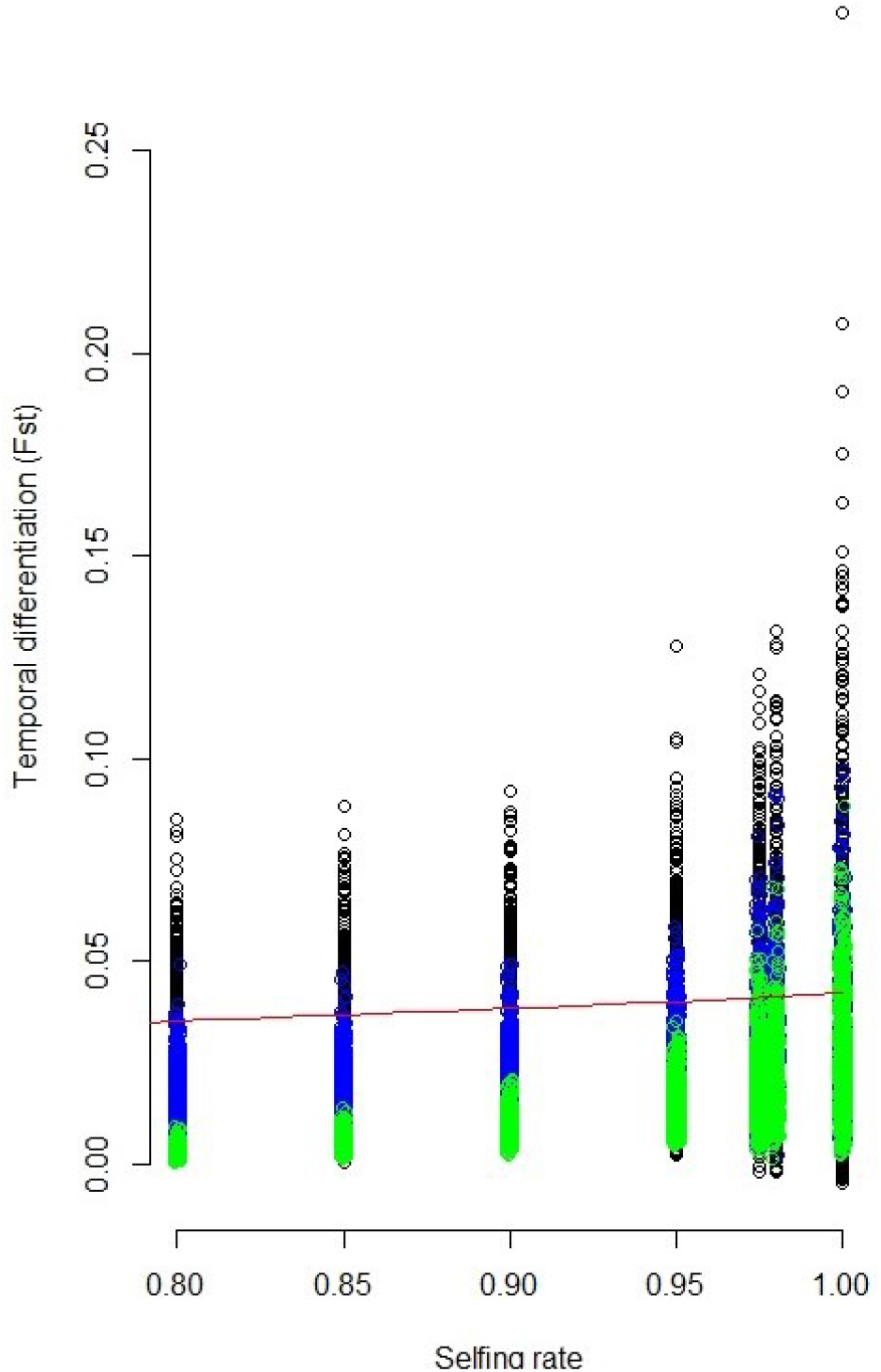
Estimates of temporal differentiation (*F_ST_*) using all loci and assuming independence (in black) or using the MLG (concatenated genotype) as alleles of a single locus, with (green) or without (blue) exclusion of the MLGs with residual heterozygosity. The red line stands for the expected value for the *F_ST_* where 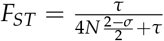 with *τ* the number of generations between the two temporal samples, *σ* the selfing rate and *N* the simulated population size.

**Figure S5.**
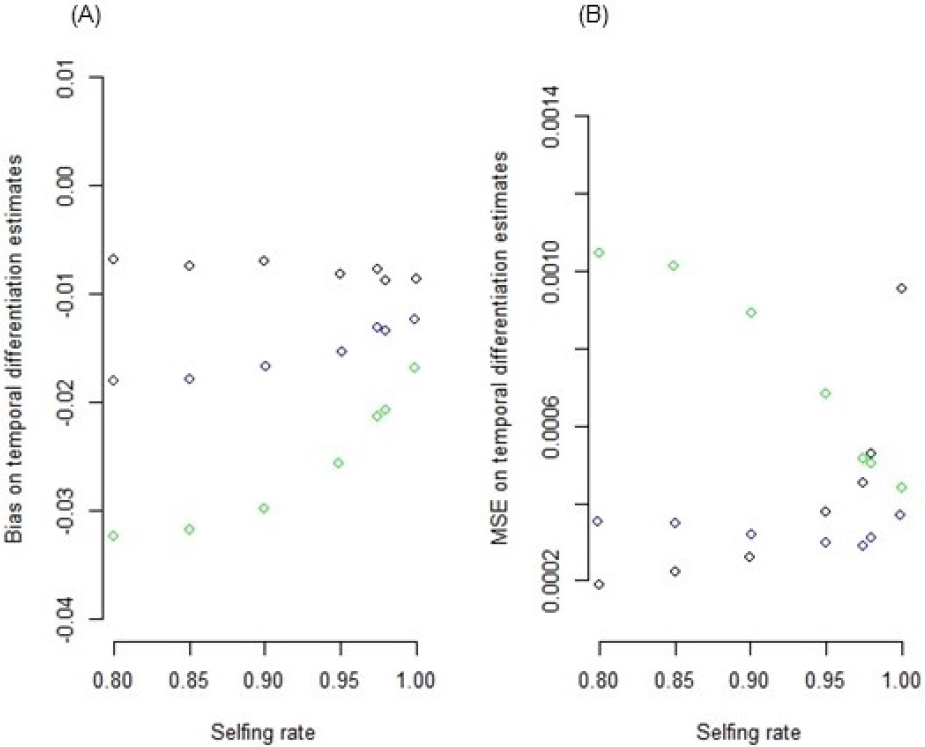
Average bias (A) and MSE (B) for the estimation of temporal differentiation (*F_ST_*) using all loci and assuming independence (in black) or using the MLG (concatenated genotype) as alleles of a single locus, with (green) or without (blue) exclusion of the MLGs with residual heterozygosity.

